# Reductive evolution of the DNA replication machinery in endosymbiotic fungi

**DOI:** 10.1101/2025.09.02.673565

**Authors:** Alexandra Dallaire, Tania Chancellor, Samik Bhattacharya, Tomoko Nishizawa, Uta Paszkowski

**Affiliations:** RIKEN CSRS, Yokohama, Kanagawa, 230-0045, Japan; Département de Biochimie, de Microbiologie et de Bio-informatique, Faculté des Sciences et de Génie, Université Laval, Quebec, G1V 0A6, Canada; Institut de Biologie Intégrative et des Systèmes, Université Laval, Quebec, G1V 0A6, Canada; Crop Science Centre, Department of Plant Sciences, University of Cambridge, Cambridge, CB3 0LE, UK; Resolve Biosciences, Creative Campus Monheim, Gebäude A03, Creative-Campus-Allee 12, 40789 Monheim am Rhein, Germany

## Abstract

The molecular machinery for replicating and repairing DNA accurately is critically important for life and highly conserved across the Tree of Life. Here we show that two major lineages of fungi, Glomeromycotina and Microsporidia, lost DNA polymerase complexes involved in replication and translesion synthesis. Catalytic and non-catalytic subunits of DNA polymerases are co-eliminated, consistent with their physical and functional interactions described in other eukaryotes. We detect lineage-specific variation in genome-wide mutation rates, showing that DNA polymerase gene losses correlate with increased genetic variation. We find that the Glomeraceae family of arbuscular mycorrhizal (AM) fungi has lived for ∼360 My without a leading strand replisome, raising the question of how these fungi can replicate DNA. We provide evidence that the cell cycle of *Rhizophagus irregularis* is active when in symbiosis with a host, but not without. This indicates a higher level of integration between AM fungi and plants than previously appreciated, and suggests the existence of a regulatory or functional contribution provided by a host to the fungal cell cycle. We propose that alternative modes of DNA replication and cell cycle provide mutational opportunities for fungal adaptation, and play roles in the evolution of endosymbioses.

## Introduction

Endosymbiotic relationships facilitated the acquisition of novel adaptive traits that propelled evolution across the Tree of Life [1-3]. Plants transitioned from freshwater to land together with intracellular filamentous fungi, assumed to be the ancestors of today’s arbuscular mycorrhizal (AM) fungi [4, 5]. The AM symbiosis has a deep evolutionary origin, and its core genetic program is conserved between plant lineages spanning 450 million years of diversification [6].

The Mucoromycota phylum comprises three subphyla (Mortierellomycotina, Mucoromycotina and Glomeromycotina), grouping species of root endophytes, plant pathogens, mycorrhizal fungi, and decomposers of plant material. Mucoromycotina and Glomeromycotina fungi are thought to have been predisposed to mutualism with plants by ancestral losses of B vitamin metabolism and plant cell wall-degrading enzymes (Figure 1a) [7]. The Glomeromycotina subphylum includes ∼300 species of AM fungi, and one of its early-diverging members, *Geosiphon pyriformis*, is the only species hosting photosynthetically active cyanobacteria [8]. All Glomeromycotina fungi carry a hallmark of obligate biotrophy, i.e. metabolic dependency caused by the loss of autonomous fatty acid biosynthesis [9-11]. Compared to Mucoromycota, the *Blastocladiomycetes* and *Chytridiomycetes* occur mainly as saprobes or parasites of plants, animals, protists or algae, and are primarily found in aquatic environments [12]. At the base of the fungal tree of life, Microsporidia are obligate intracellular parasites of animals, and their close relatives Cryptomycota are another group of intracellular parasites of algae, aquatic fungi (e.g. *Rozella*), crustaceans (e.g. *Mitospodidium*), and amoebae (e.g. *Paramicrosporidium*).

**Figure 1.**
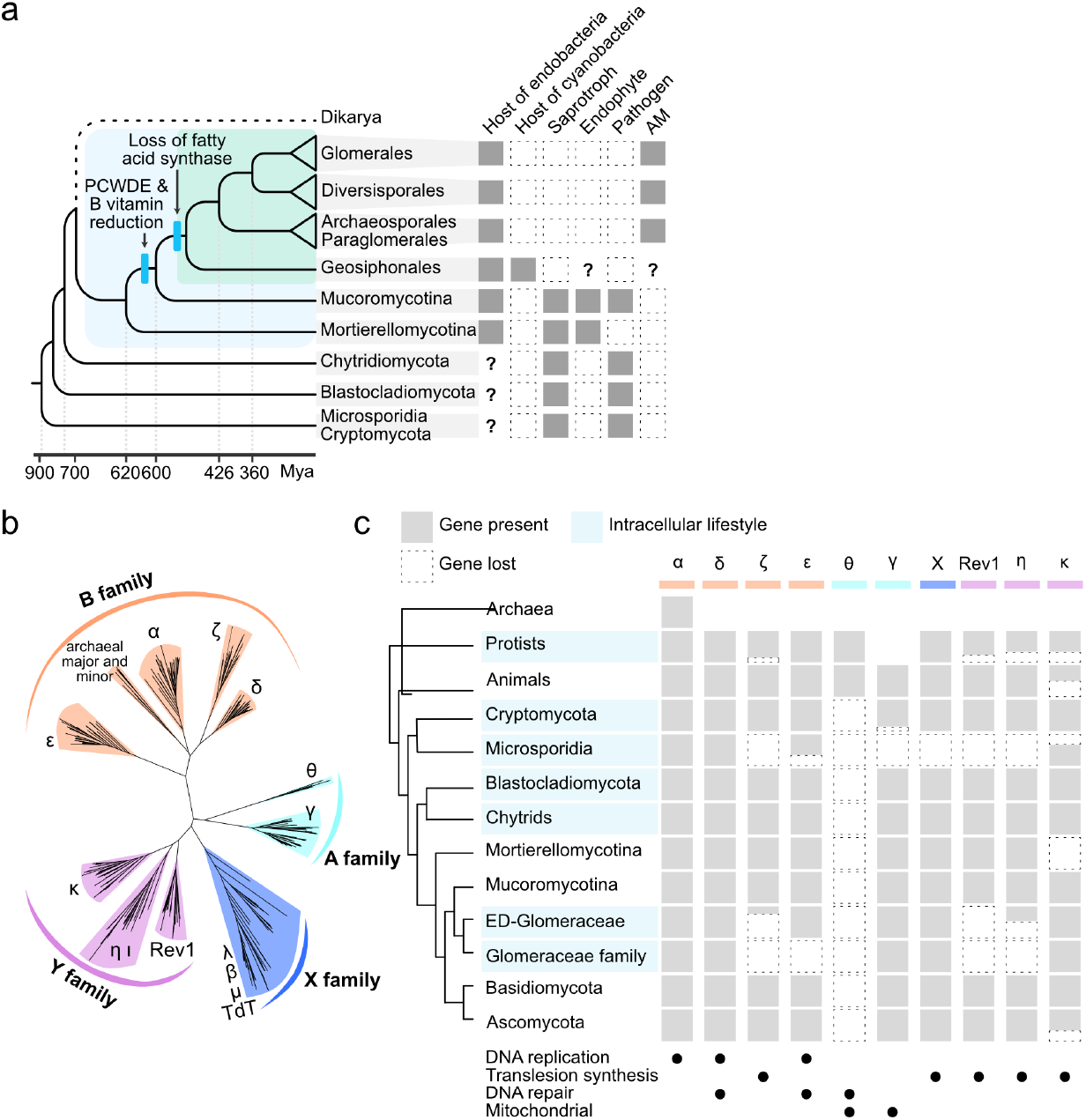
Traits and phylogenetic analysis of fungal DNA polymerases. a) Trait presence and absence are indicated by full and dotted-line squares respectively, and lack of data by a question mark. The Mucoromycota clade is delineated in blue, and Glomeromycotina in green. PCWDEs = plant cell wall-degrading enzymes; AM = arbuscular mycorrhiza. Approximate tree dating based on [38] and traits based on [15-17]. b) Phylogeny of the A, B, X and Y family DNA polymerases of fungi, animals, protists and archaea. ML phylogeny built with FastTree, bootstrap=1000. c) Summary of gene presence (grey box) and loss (dotted empty box). Variation within species of the taxon analysed is represented by proportionally shaded/dotted boxes. Taxa with intracellular lifestyles are highlighted by blue rectangles. The origin of polymerase gamma (Polγ) is uncertain, therefore no box is shown in protists. Black dots indicate gene function as inferred from experimental work in yeast and animal cells. ED = early-diverging.

Lifestyle transitions from free-living to obligate endosymbiosis in prokaryotes are thought to occur in phases, where genome expansion, driven by the acquisition of mobile elements, precedes genome reduction [13]. Long-term endosymbiotic bacteria display small genome sizes, substantial gene loss, and reduced DNA replication and repair pathways [14-19]. Independent pervasive losses of replicative DNA polymerases (DnaE^Polα^, DnaG^Prim1/2^, DnaQ^Polε^), DNA clamp (DnaN^PCNA; Rad9/Rad1/Hus1^), mismatch repair (MutL^Mlh1/2/3/Pms1^, MutS^Msh2/3/6^), base excision repair and homologous recombination factors (RecA/B/C/D^Rad51^), can contribute to enhanced mutation rates and genome instability [20-23], but can also provide opportunity for adaption [24]. In eukaryotic microbes, reductive evolution of the mismatch repair pathway has been reported in multiple lineages of pathogenic fungi, and was proposed to drive mutator phenotypes for rapid adaptation to plant and animals hosts [25-30].

Here, we investigate the completeness of the DNA replication machinery in lineages of endosymbiotic fungi. Through evolutionary analysis of replisome components, we reveal lineage-specific losses of DNA polymerase genes and associated co-factors in obligate mutualists of the Glomeromycotina subphylum and in obligate parasites of the Microsporidia phylum. Questioning the evolutionary and mechanistic impacts of replisome reduction, we show that polymerase gene loss is associated with higher intraspecific mutation rates, and with the co-elimination of replisome components. Replisome reduction may therefore cause lineage-specific variation in evolutionary rates, and is a trajectory shared between eukaryotic and prokaryotic symbionts with an obligate intracellular lifestyle. Species of the Glomeraceae family of AM fungi persisted over 360 My with the most reduced replisome, and likely coordinate their cell cycle with factors provided by symbiosis with a host.

## Results

### Loss of DNA polymerase genes in eukaryotic intracellular parasites and mutualists

DNA polymerases are classified into seven major families (A, B, C, D, E, X and Y) according to their sequence homology (Table 1) [31]. The C, D and E families are exclusive to prokaryotes and are not analysed further in this work. A-family polymerases replicate nuclear and organellar genomes, and animal Polθ has additional roles in alternative non-homologous end-joining and recombination [32, 33]. The B-family of replicative DNA polymerases includes the DNA primase alpha (Polα) and the leading and lagging strand polymerases epsilon (Polε) and delta (Polδ). Pol zeta (Polζ) is the only B-family polymerase playing a critical role in translesion synthesis, a mutagenic mechanism that allows recovery of replication forks at stalled sites. X-family polymerases are error-prone and function in base excision repair (BER) and non-homologous end joining (NHEJ) [34]. Polymerases of the Y-family (Rev1, Polη, Polι, Polκ) are all involved in translesion synthesis and somatic hypermutation, and typically have low processivity and high error-rates [35].

**Table 1:**
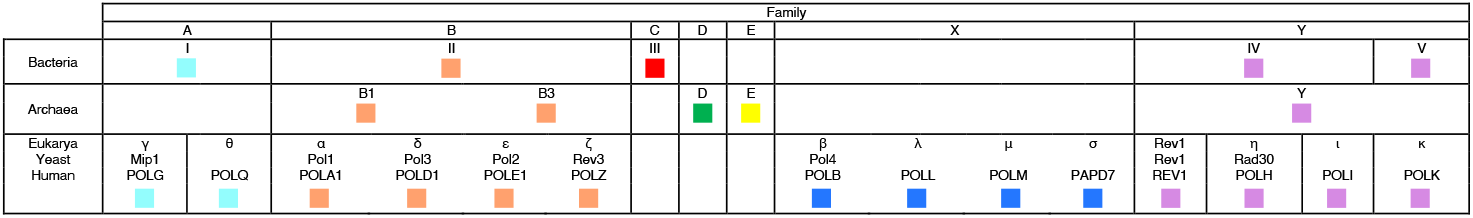
DNA polymerase family nomenclature in bacteria, archaea, yeast and human.

Using a phylogenomics framework and homology inference, we identified and classified the DNA polymerases of representative fungi, animals, protists and archaea (Figure 1b, Table S1, S2). We found that Polζ was lost in the intracellular Microsporidia and Glomeromycotina species, with the notable exception of *Geosiphon pyriformis* (Figure 1c, Table S3). The A-family polymerase Polθ is found in protists and animals, but not fungi, whereas Polγ is found in animals and fungi, but not protists (Figure 1c, S1ab and Table S3). Therefore, Polθ was present in the last eukaryotic common ancestor and was lost in fungi, while Polγ originated more recently and was present in the ancestor of Opisthokonta (Figure S2c). Polγ polymerases are thought to derive from T7-like PolA (not Polθ) [32], suggesting that this mitochondrial replication factor was acquired from a bacteriophage. Microsporidia lost all A-family polymerases, and while they have long been thought to be amitochondriates, they do carry cryptic mitosomes which lack a genome and the associated DNA replication machinery [36]. We confirm previously reported losses of Polκ in *D. melanogaster* and *S. cerevisiae* [37], and report new examples of Rev1, Polκ and Polη losses in protists, Cryptomycota, Mortierellomycotina and Microsporidia (Figure 1c, Figure S1 and Table S3). All Glomeromycotina species have lost Rev1, and partial losses of Polζ, Polη and Polε are observed throughout the subphylum. Homologs determined based on protein structure could not be identified (see Materials and Methods), and we detected no significant relationship between the number of polymerase complex genes and the proportion of benchmarking universally conserved single-copy ortholog (BUSCO) genes in Microsporidia and Glomeromycotina, supporting true biological variation rather than artifacts due to incomplete genomic data (Figure S2ab). Analysis of bitscores obtained through pairwise sequence alignments show that Polζ, Polη and Polε are above detectability threshold in the Mucoromycota phylum and are not diverging radically in Glomeromycotina species (Figure S2c). Gene loss therefore likely explains our observations, unless clade-specific accelerations in divergence led to both sequence- and structure-level homology detection failure. We conclude that DNA polymerases with function in replication, translesion synthesis and DNA repair can be lost in eukaryotic microbes, predominantly in intracellular symbionts.

### Lineage-specific loss of DNA polymerases in Glomeromycotina and Microsporidia

Sequenced genomes of the Glomeraceae family of AM fungi are missing four out of the nine DNA polymerases which are otherwise broadly conserved in free-living eukaryotes. An unexpected loss is the leading strand DNA polymerase Polε, an enzyme essential for DNA replication, cell cycle progression, genome stability and epigenetic regulation in human and yeast [21]. Surveying gene content at fine taxonomic scale revealed that the catalytic subunits of polymerase complexes were lost sequentially: Rev1 was first lost in the ancestor of Glomeromycotina, followed by Polζ, Polη and Polε at each of the next phylogenetic splits (Figure 2). Remarkably, these gradual losses match the speciation pattern, suggesting that altered replication and associated changes in mutation rates played a role in the divergence of ancient populations [39].

**Figure 2.**
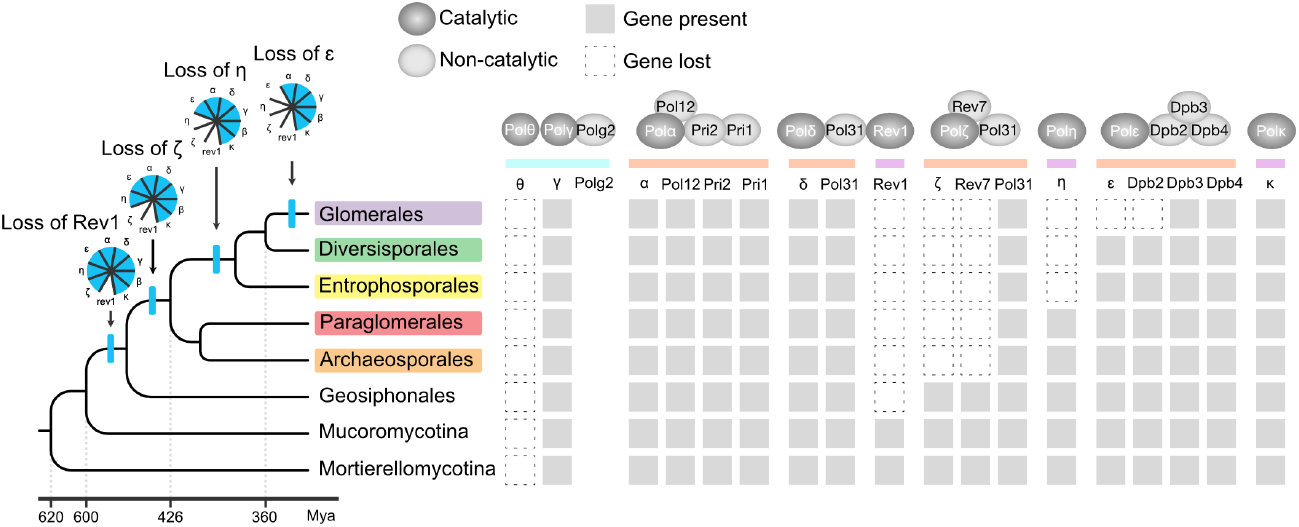
Lineage-specific losses of catalytic and non-catalytic subunits of DNA polymerases in Mucoromycota. Gene presence (grey box), gene loss (dotted white box), and non-conserved genes (no box). Dating of Mucoromycota tree from [38].

DNA polymerases α, δ, ε, ζ and γ function as holoenzymes through physical interaction with non-catalytic accessory subunits [40]. We assessed the presence of holoenzyme subunits in Glomeromycotina species and found that the non-catalytic subunits Rev7 and Dpb2 were lost concomitantly with their catalytic partners (Figure 2, Figure S3). This co-elimination pattern likely reflects the physical and functional interactions between the subunits [41]. The remaining non-catalytic subunits Pol31 and Dpb3/4 are respectively shared with the Polδ holoenzyme and with chromatin remodelling complexes [40, 42, 43]. Their retention suggests selective pressure to maintain alternative functions.

Since Glomeromycotina and Microsporidia species similarly lack Rev1, Polζ, Polη and Polε, we wondered if these genes were lost in the same sequential order in both lineages. In a finer-scale analysis, we identified DNA polymerases in 28 Microsporidia species and in all available genomes of their early-diverging sister lineage Cryptomycota (also called Rozellomycota or Rozellida). While the mitochondrial Polθ is absent in fungi, we observe that Polγ was lost in the transition between Cryptomycota and Microsporidia, together with the mitochondria (Figure S4)[44]. Polg2, the processivity factor of Polγ in animals, originated in Metazoa and was lost independently in the Helminthes and Annelida phyla, in addition to the previously reported loss in Nematoda (Figure S5, Table S4, Table S5) [45, 46]. In the transition between Cryptomycota and Microsporidia, Polζ was lost first, followed by Rev1 and Polη (Figure S4). Translesion polymerase complex reduction therefore precedes the loss of the mitochondria and clade-specific Polε and Polκ losses in the Microsporidia lineage. Like in Glomeromycotina, Microsporidian Polζ and Polε were co-eliminated together with their accessory subunits Rev7 and Dpb2 (Figure 2, Figure S4). While new Cryptomycota genomes will be required to retrace the sequence of Rev1 and Polη losses, our analysis indicates that in both Microsporidia and Glomeromycotina, translesion polymerases Polζ, Rev1 and Polη were lost before the mitochondria, and before Polε.

Loss of a replicative DNA polymerase is rare in eukaryotes, and co-elimination of Polε and Dpb2 in the Glomeraceae family suggests the existence of a substantially remodelled replisome. We next sought to identify core replisome factors in Glomeromycotina, relying on direct protein interactions characterised by cryo-EM in yeast and human [47-49] (Figure 3a). We found that the alternative clamp loader Ctf18-RFC, which specifically functions with the leading strand replication machinery via an interaction with Polϵ [50, 51], is missing three factors, Ctf18, Ctf8 and Dcc1 in Glomeraceae species (Figure 3bc). Thus, additional direct interactors of Polε that are not shared with Polδ or another complex are co-eliminated in the Glomeraceae family.

**Figure 3.**
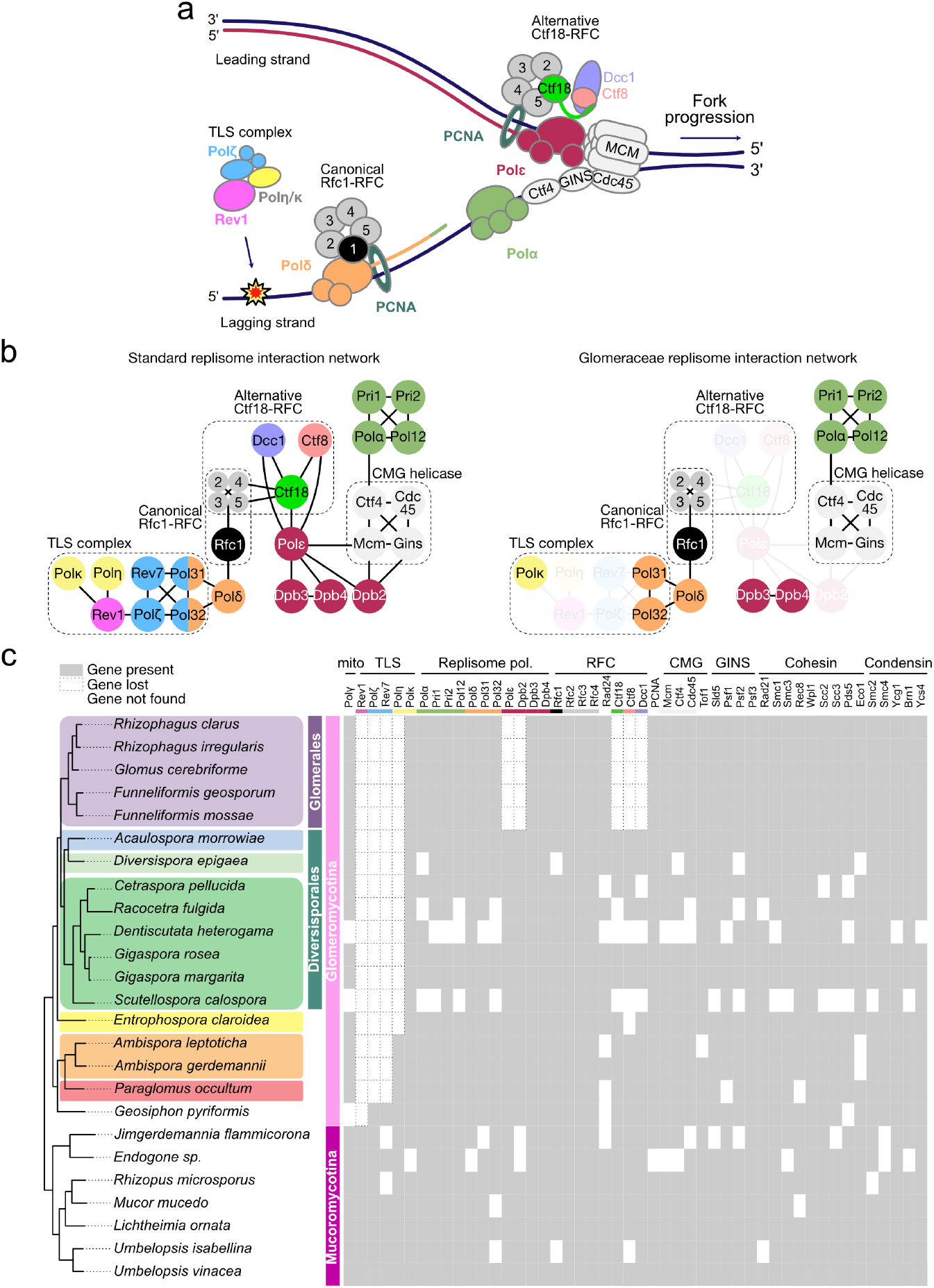
Loss of Polε-associated replisome factors in the Glomeraceae family. a) Model of a standard replisome, with the canonical Rfc1-RFC clamp-loader associated with Polδ, and the alternative Ctf18-RFC clamp-loader associated with Polε. b) Summary of direct interactions between replisome factors characterised by cryo-EM in human and yeast (left), and proposed Glomeraceae replisome (right). c) Replisome factor presence and absence in Mucoromycota. Species missing several genes, observed as horizontal white lines, were considered to have incomplete genome assemblies or annotations, in which case missing genes were instead labelled ‘not found’.

Because the loss of function of replicative or translesion DNA polymerases can contribute to enhanced mutation rates and genome instability, we quantified inter-strain genetic variation in Glomeromycotina and Microsporidia species with different polymerase repertoires. In Glomeromycotina, strains of Polη-; Polε-*R. irregularis* (Glomeraceae) and Polη-*C. pellucida* (Gigasporaceae) have lower whole-genome alignment rates and higher single nucleotide polymorphisms (SNP) rates than the Polη+; Polε+ species *Entrophospora candida* (Entrophosporaceae) (Figure 4). In Microsporidia, Polκ-species display higher inter-strain SNP rates than the Polκ+ *Vairimorpha ceranae*. The loss of Polη and Polε in Glomeromycotina and of Polκ in Nosematida (Microsporidia) is therefore associated with greater genetic diversity.

**Figure 4.**
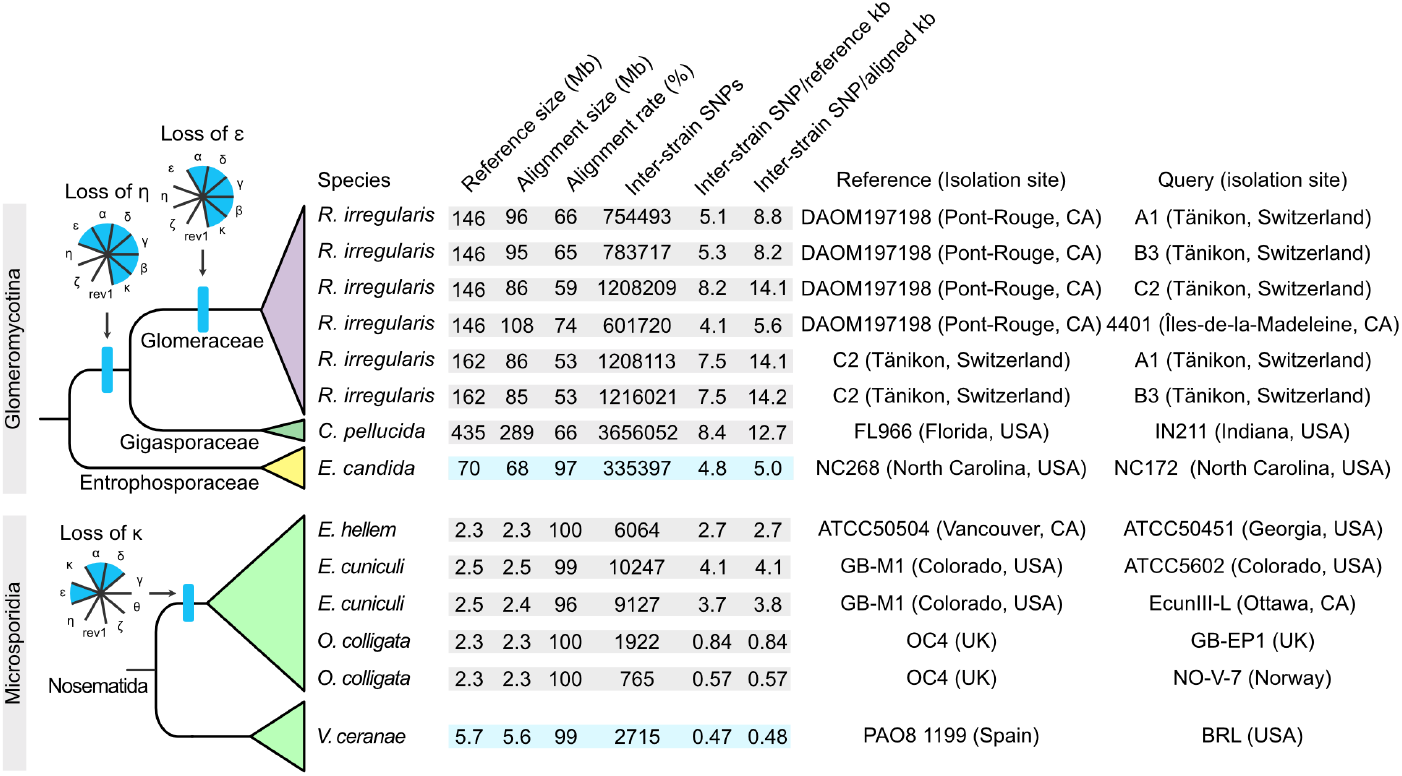
Inter-strain genomic variation in Glomeromycotina and Microsporidia with different DNA polymerase repertoires. Whole-genome alignment rates and SNP rates are reported for pairwise comparisons of Glomeromycotina strains from species with or without Polη and Polε, and of Microsporidia strains from species with or without Polκ.

Together, our data shows that similarly to several groups of endosymbiotic bacteria, the Glomeromycotina and Microsporidia lineages experienced reductive evolution of polymerase complexes typically involved in DNA replication, translesion synthesis and DNA repair. We uncover variable losses across AM fungi, with the model family displaying the most reduced replisome. We provide evidence that loss of polymerase genes is associated with increased genetic variation.

### Activity of the *R. irregularis* cell cycle when in symbiosis with a host, but not without

The model species of AM fungi, *R. irregularis*, is part of the Glomeraceae family which lacks a leading strand replisome. We hypothesized that reduction of this machinery causes dependence to a regulatory or functional contribution provided by symbiosis with a host. To test whether DNA was synthesised by *R. irregularis* in the absence of a host, we germinated spores aseptically *in vitro*, and quantified the amount of chromosomal and mitochondrial DNA over three weeks. While DNA content increases during growth of filamentous fungi [52], we detected no significant increase of the amount of *R. irregularis* chromosomes over time, indicating that while the fungus grows and branches, its cell cycle is likely inactive (Figure 5a). We quantified gene expression during spore germination compared to colonized roots, and found that most DNA replication and repair genes are lowly expressed in absence of a host, and strongly up-regulated *in planta*, coinciding with the activation of fatty acid biosynthesis in response to host provision (Figure 5b). Genes down-regulated in roots include subunits of the Cohesin complex Smc1 and Smc3, and up-regulated genes include all subunit of the Condensin I complex (Smc2, Smc4, Brn1, Ycs4 and Ycg1). Cohesin activity stops at the end of the G2 phase, while Condensin I functions during anaphase and telophase in the animal cell cycle [53]. Therefore, our data suggests that nuclei in asymbiotic spores are in the G1 phase of the cell cycle, and that colonised roots are enriched in mitotic nuclei. Since anaphase and telophase are characterized by the separation of sister chromatids which are pulled towards opposite poles, we sought to visualise mitotic fungal chromosomes in presence and absence of a host. In colonised roots, we identified pairs of regularly spaced nuclei appearing to undergo anaphase or telophase in fungal vesicles, structures historically thought to serve for lipid storage (Figure 5c). In spores germinated without a host, paired nuclei can be found in hyphae, albeit at a lower rate, and may either be dividing or aggregating randomly (Figure 6d,e). Together, this indicates that the *R. irregularis* cell cycle is active when in symbiosis with a host, and may specifically be localised to vesicles.

**Figure 5.**
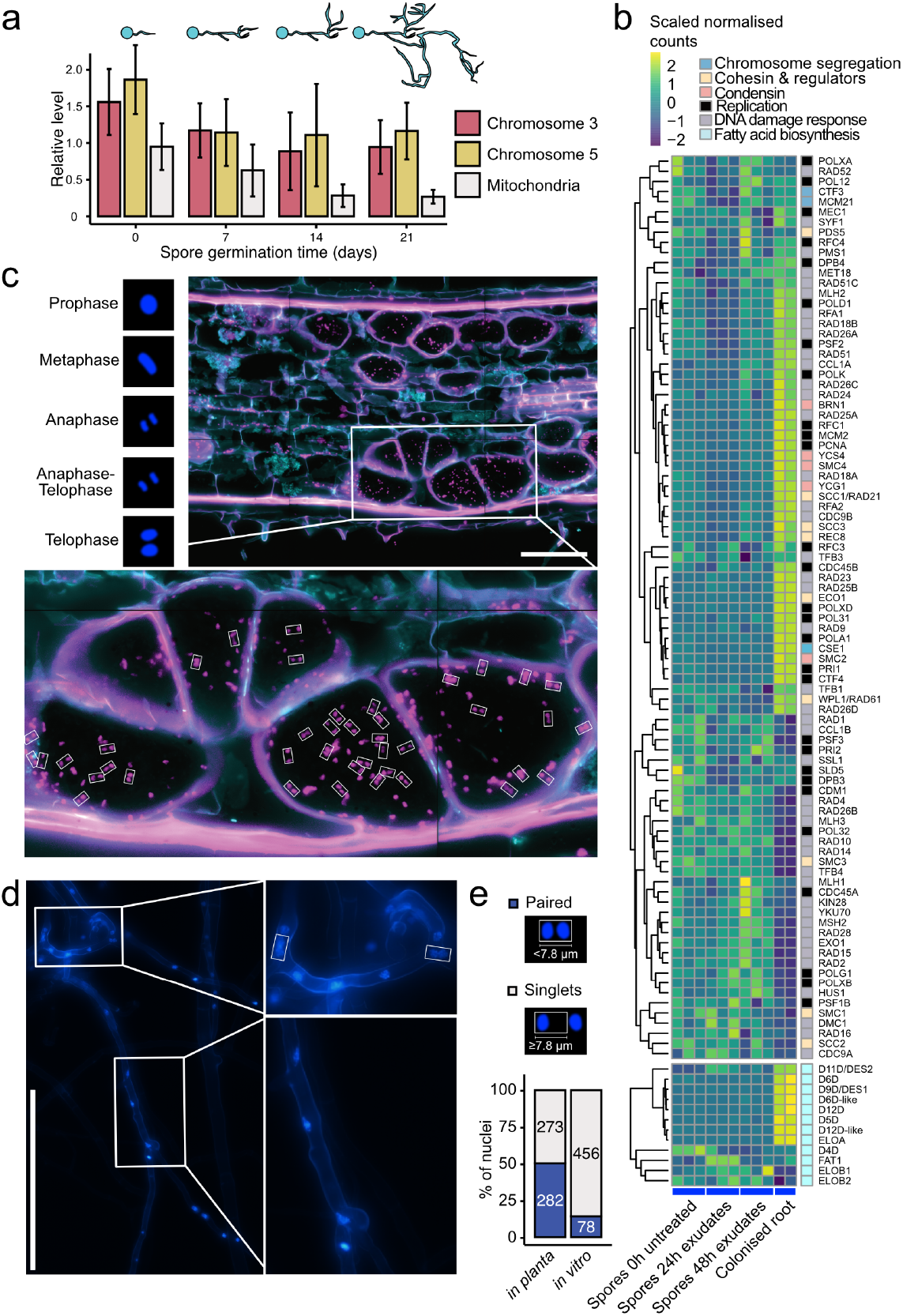
DNA replication of Rhizophagus irregularis with and without a host. a) Nuclear and mitochondrial DNA measured by qPCR and normalised on the extraction spike-in pUC19 following spore germination in water for 21 days at 28^°C^. The mean and standard deviation are plotted for 5 replicates per timepoint. A schematic illustration shows spore germination and branching. b) Heatmap and hierarchical clustering of scaled normalised counts of replication and fatty acid biosynthesis genes during spore germination and in planta. Four conditions were used, a 0-h no treatment, 24-h rice exudate treatments, 48-h rice exudate treatments (three replicates per treatment), and colonized roots (2 replicates). c) Composite confocal microscopy image of a rice root colonised by R. irregularis, stained with DAPI (magenta) and wheat germ agglutinin conjugated to Alexa Fluor 488 (cyan). The inset shows fungal vesicles with regularly spaced nuclei appearing to undergo anaphase or telophase. A schematic diagram shows expected chromosome condensation and separation during mitosis. Bar 100 μm; small white boxes 7.8×4.2 µm. d) Maximum projection fluorescence microscopy of 7-day in vitro germinated spores (host-free), stained with DAPI (blue). Bar 100 μm; small white boxes 7.8×4.2 µm. e) Stacked bar graphs showing the proportion of paired and singlet nuclei in planta (vesicles shown in c) and in vitro (hyphae shown in d). Total counts are shown inside the stacks.

## Discussion

Given the estimated divergence time between Glomeromycotina and Mucoromycotina, our work indicates that in AM fungi, DNA polymerase loss evolved gradually, over at least 500 My. For up to 360 My, the Glomeraceae family lived with a most reduced polymerase repertoire, and achieved DNA replication without the leading strand DNA polymerase Polε. In *S. cerevisiae* and *S. pombe*, Polδ can function on both strands and proofreads errors made by Polε, although this results in increased mutagenesis [54, 55]. This form of redundancy could enable leading strand synthesis in the Glomeromycotina species that have lost Polε, and suggests the existence of a mechanism that is distinct from currently known modes of replication. This implies consequences on the function of multiple replisome factors that physically interact with Polε [47-49]. We find that Polε was co-eliminated together with its non-catalytic subunit Dpb2, and the alternative clamp-loader factors Ctf18, Ctf8 and Dcc1 [56]. In addition to its replicative role, the Polε-bound Ctf18-RFC is important for sister chromatid cohesion and signals fork stalling to activate the S-phase checkpoint [57]. The Ctf18 gene was discovered in a screen for yeast mutants with elevated rates of mitotic chromosome loss [58], and its loss of function increases levels of mitotic recombination (also called parasex), a repair process associated with increased mutagenic recombination [59]. The replisome of Glomeraceae species therefore appears to be substantially re-modelled as it has no leading strand DNA polymerase and no leading strand clamp loader. This gene loss pattern is likely not stochastic, and can either be adaptive, or caused by relaxation of constraints to maintain obsolete functions. Dependence on a host for DNA replication has so far only been reported in viruses, and may be prohibited by structural barriers between plants and AM fungi. Extracellular vesicles forming at the arbuscular interface [60] could however provide opportunity for hijacking the host replication machinery. A “use it or lose it” principle is hardly applicable to the DNA replication machinery, and rather plausibly reflects a scenario of adaptive evolution in which the loss of function is advantageous and positively selected.

In human and yeast, the loss of function of Rev1, Polζ, Polη or Polε causes genomic instability and cell cycle defects [20, 61-66]. In the absence of TLS polymerases, damaged regions are resolved using mitotic recombination [67]. It is therefore conceivable that DNA polymerase loss in AM fungi and Microsporidia results in lineage-specific variation in replication accuracy, mutation rates and chromosomal rearrangements. Consistent with this idea, we find that loss of DNA polymerases is associated with lower inter-strain genome alignment rates and higher SNP rates in specific Glomeromycotina and Microsporidia lineages. We propose that the loss of these genes could be an underlying cause of the exceptionally large pangenome (up to 50%), high inter-strain variability in chromosome lengths and genome reshuffling reported in *R. irregularis* [68, 69]. New genome assemblies will be necessary to draw conclusions on differential genetic variability across all families of Glomeromycotina and Microsporidia.

Loss of core complexes involved in maintenance of genetic information is rare in eukaryotes but has been described in microorganisms which abandon their free-living lifestyle to become intracellular symbionts. In Microsporidia and bacterial endosymbionts of insects, genome decay via accumulation of deleterious mutations and gene loss primarily stem from genetic drift in small, genetically bottle-necked populations of vertically transmitted endosymbionts [14-19]. Conversely, AM fungi are endemic in terrestrial ecosystems, have a wide host range and are horizontally transmitted. Despite metabolic dependence on their hosts, AM fungi are unlikely to experience population bottlenecks, and their large population size conceivably increases the efficiency of purifying selection. An additional level of selection can act on genome ‘populations’ within their massively multi-nucleate mycelia. Therefore, even though AM fungi have lost genes involved in DNA replication, they may not follow the drifting path of Microsporidia towards extreme genome reduction and compaction.

In Microsporidia, the Glugeida species that have lost Polε form a clade including the highest number of hyperparasites and species able to infect more than two host groups [70]. In the Glomeraceae family, loss of Polε and replisome reduction coincide with this lineage having the highest speciation rate and the largest ecological niche width among Glomeromycotina [39]. Early-diverging Ambisporaceae and Paraglomeraceae were shown to provide little to no benefit to plants, compared to late-diverging Diversisporaceae, Entrophosporaceae and Glomeraceae [71]. Therefore, the number of DNA polymerases in AM fungi appears to inversely correlate with 1) the ability to stimulate plant growth and nutrient uptake, and 2) speciation rates and range expansion. Because the loss of vitamin biosynthesis and cell wall-degrading enzymes is shared between Mucoromycotina and Glomeromycotina, and given the unknown mycorrhizal status of Geosiphonales, the lack of Polζ, Polη and Polε currently represent more accurate genomic indicators of AM symbiosis than the loss of fatty acid synthase alone.

Because of their obligate biotrophy relying on fatty acids supplied by plants, host-free *in vitro* culture of *R. irregularis* results in hyphal branching without lifecycle completion to a next generation. An aseptic culture system relying on the provision of myristate and palmitate allows limited growth of one or two generations of secondary spores, which are however smaller than those generated symbiotically [72]. Our findings suggest that aseptic culture of *R. irregularis* may be limited by DNA replication, and prompts to examine the connection between the fungal nutritional status and cell cycle activity. In addition, since homologous recombination can only happen in the G2 phase of the cell cycle [73], we propose that genome editing approaches for Glomeraceae species may be more efficient when targeted *in planta*, rather than in asymbiotic spores.

Our findings have implications for understanding the evolution of obligate endosymbionts. Loss of components of DNA replication and repair systems is a reductive trajectory shared by prokaryotic and eukaryotic obligate endosymbionts. Population genomics, synthetic biology and transformation efforts deployed to study endosymbiotic systems may consider the evolvability of DNA replication fidelity and mutation rates, as well as alternative modes of cell cycle regulation and DNA repair.

## Supplemental Figures

**Figure S1.**
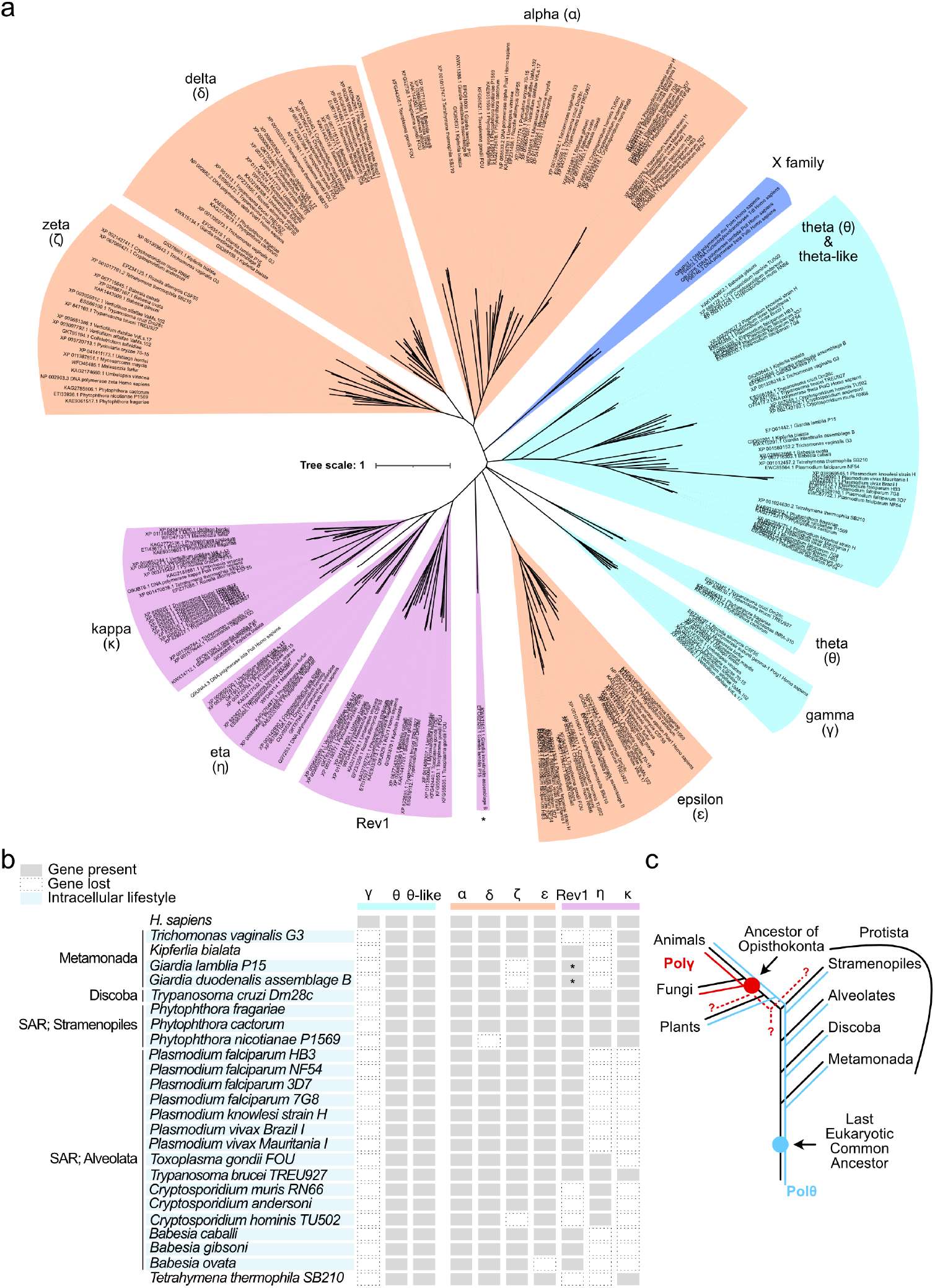
Phylogenetic analysis of protist DNA polymerases. a) Phylogeny of the A, B, X and Y family DNA polymerases of selected protists and outgroups. Giardia Rev1-like proteins are marked by a star (*). ML phylogeny built with FastTree, bootstrap=1000. b) Summary of gene presence (grey box) and loss (dotted empty box). Species with intracellular lifestyles are highlighted by blue rectangles. c) Proposed evolutionary history of mitochondrial Polθ (blue) and Polγ (red) in the Eukaryotic Tree of Life (black).

**Figure S2.**
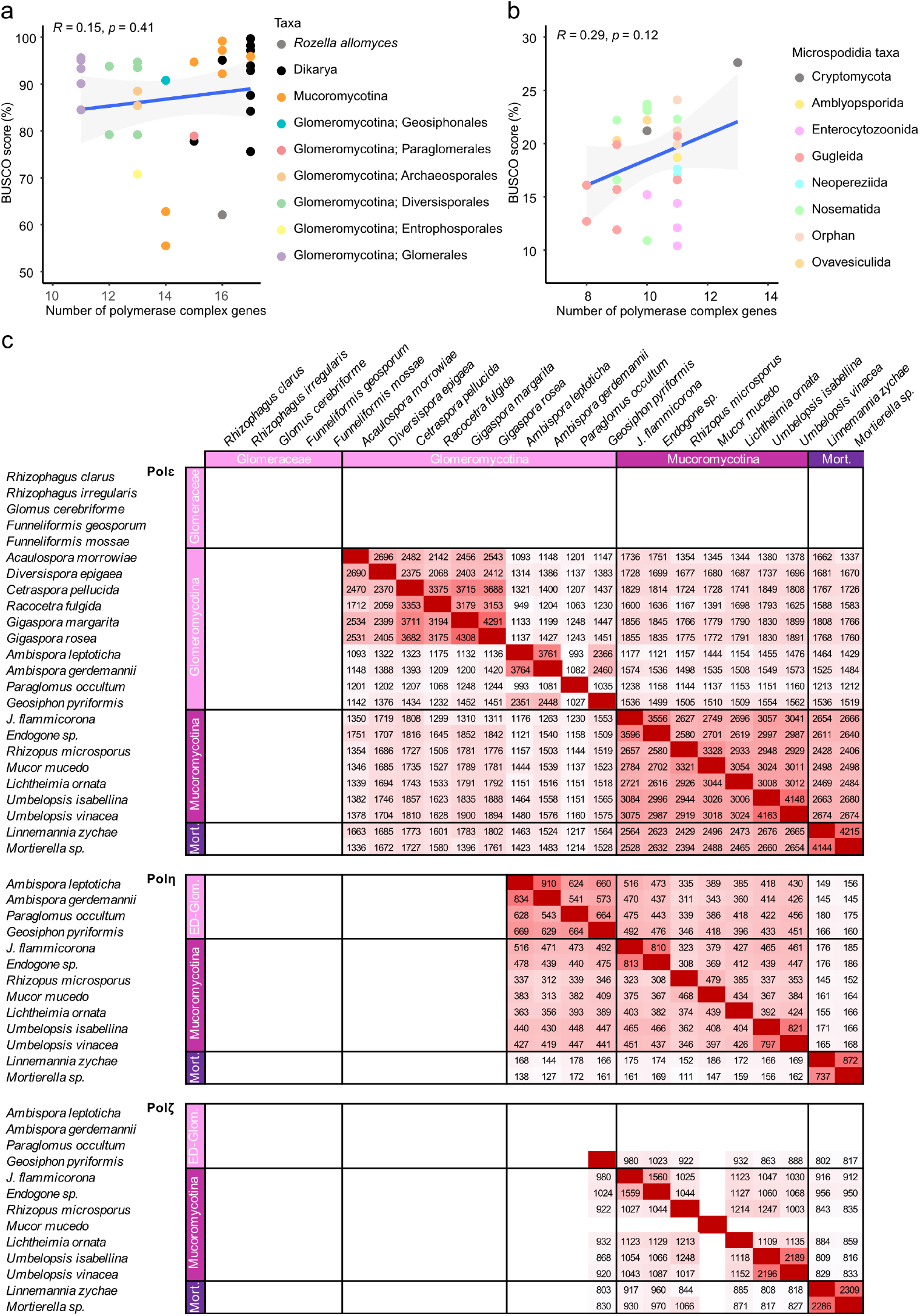
Assessment of assembly completeness and homology detection limits. Relationship between the number of polymerase complex genes (Table S3) and proportion of BUSCO genes detected in A) Mucoromycota, Dikarya and Rozella, and B) Cryptomycota and Microsporidia species. C) Bitscores from pairwise sequence alignments of Mucoromycota Polε, Polη and Polζ, visualised by individually scaled heatmaps.

**Figure S3.**
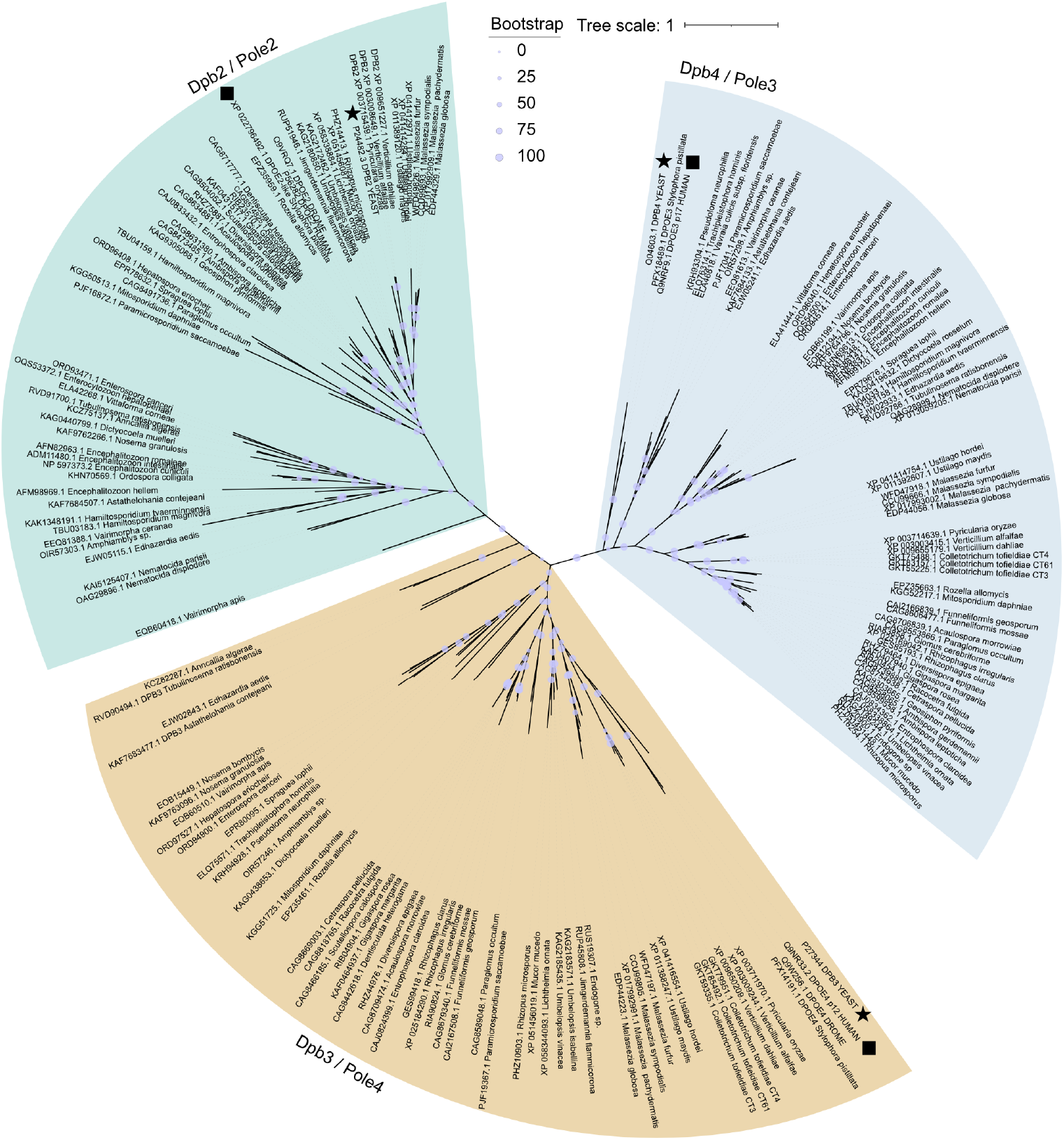
Classification of non-catalytic subunits of Polε. Unrooted phylogenetic tree of DNA polymerase delta subunits Dpb2/Pole2, Dpb3/Pole4 and Dpb4/Pole3. Dpb2 is the largest subunit (average = 451 aa), followed by Dpb4 (average = 180 aa) and Dpb3 (average = 177 aa). *Saccharomyces cerevisiae* ✶, *Homo sapiens* ◼. Bootstrap is shown as a percentage of 1000 replicates.

**Figure S4.**
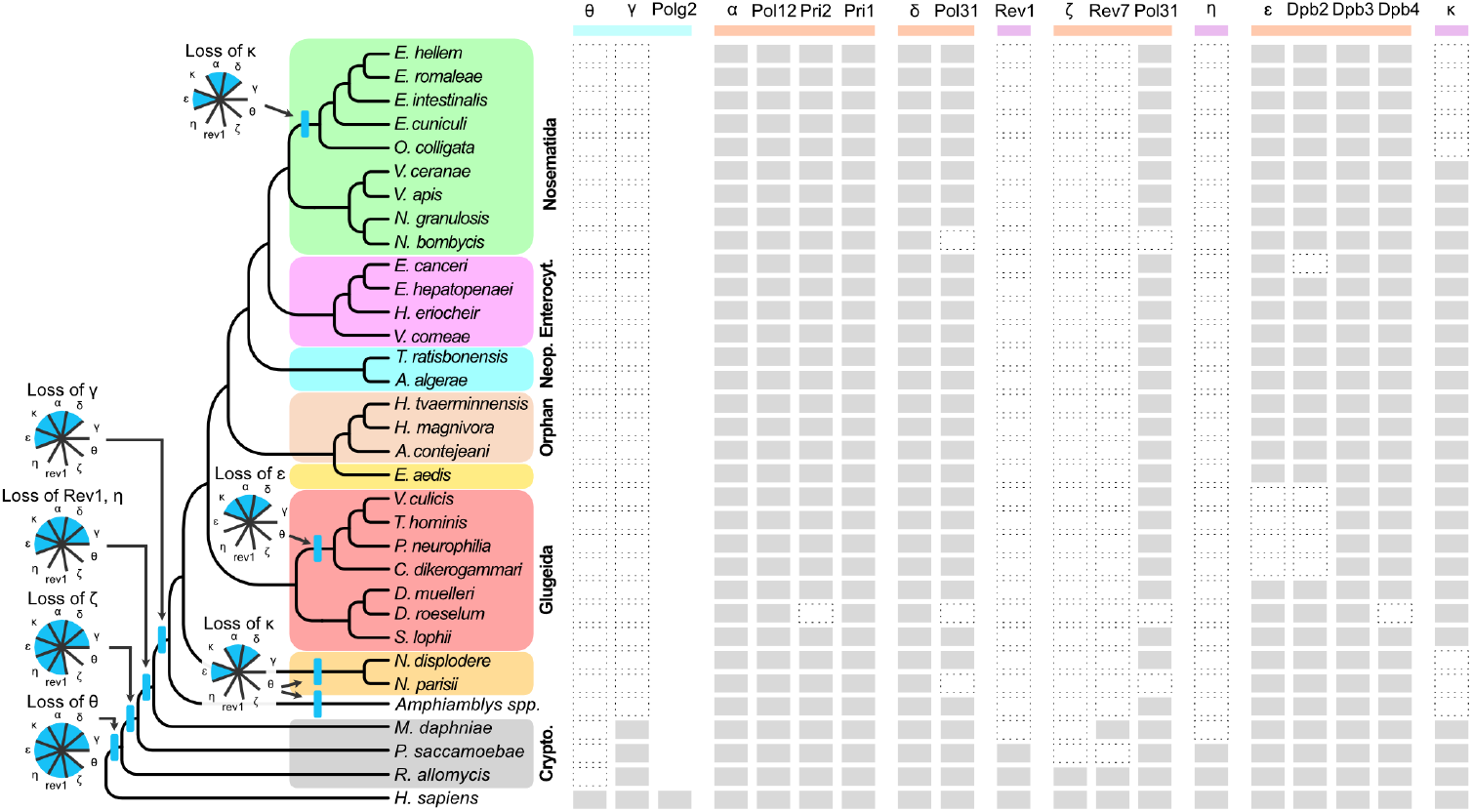
Lineage-specific losses of catalytic and non-catalytic subunits of DNA polymerases in Microsporidia. Gene presence (grey box), gene loss (dotted white box), and non-conserved genes (no box). Genome-level phylogeny of Microsporidia was reproduced from [70].

**Figure S5.**
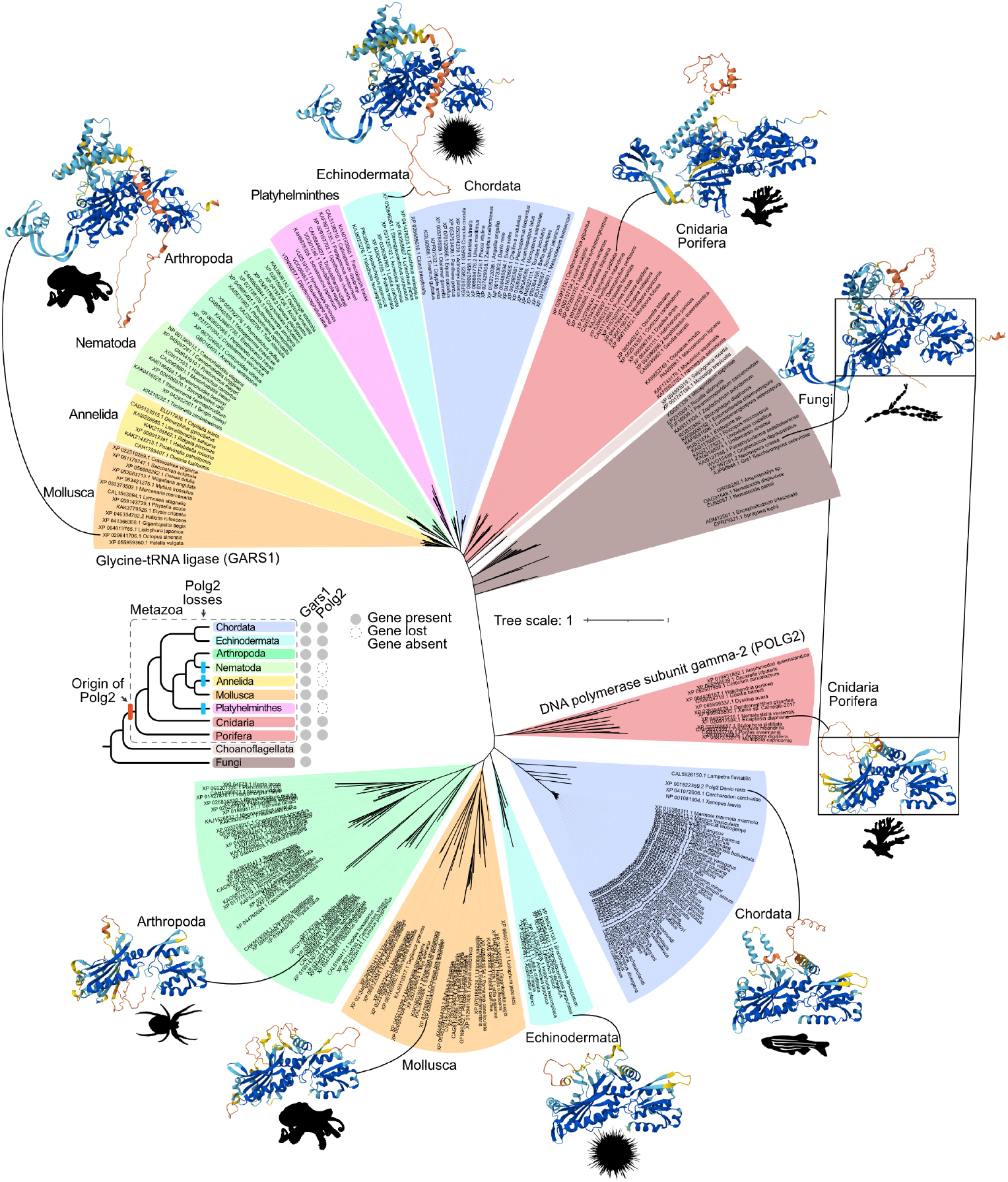
Origin and losses of Polg2 in Metazoa. Unrooted phylogenetic tree of DNA polymerase subunit gamma-2 (POLG2) and related glycine-tRNA ligase (GARS1) in Opisthokonta, with protein structures displayed for selected species. The proteins most closely related to POLG2 outside of Metazoa are GARS1. POLG2 and GARS1 share a rare anticodon-binding domain fold ([74], highlighted by linked black boxes), but are differentiated by their sequences, average lengths (416 and 718 amino acids respectively), and overall structures. The inset panel summarizes gene presence and loss (grey and white circles respectively), showing that Polg2 is absent in Fungi and Choanoflagellata, and therefore originated in the ancestor of Metazoa (red bar). Polg2 was independently lost in the Platyhelminthes, Annelida and Nematoda phyla (blue bars).

## Materials and Methods

### Comparative genomics

For comparative genomic analyses, we downloaded the genome assemblies and proteomes of 69 fungal species from NCBI (Table S1). BUSCO v5.8.2 was run on all assemblies using the fungi_odb10 database (Table S1). Orthologous groups within the Mucoromycota and Dikarya were inferred using OrthoFinder 2.5.5 [75] with default parameters. Yeast and human proteins were used to identify and annotate orthogroups corresponding to known DNA replication and repair factors (Tables S2-S5). Successive protein sequence searches for homology inference were performed using DIAMOND [76], PSI-BLAST [77] and TBLASTN [78]. Structural similarity searches were done by modelling proteins with AlphaFold3 [79], and using FoldSeek [80] with the iterative search option and appropriate taxonomic filters (e.g. Glomeromycotina). Protein sequences were aligned with MAFFT [81] with default parameters, phylogenetic trees were built using IQ-Tree 2.4.0 [82] or FastTree [83] (as described in figure legends), and trees were visualised in iTOL [84].

### Whole-genome SNP analysis

Genome-wide SNPs were measured in a pair-wise manner in Glomeromycotina and Microsporidia strains (accessions listed in Figure 5 and Table S1) using PhaME [85]. The MUMMER delta-filter was set at a minimum alignment uniqueness of 30% (-u 30).

### Gene expression analysis

Publicly available RNA-seq data (accessions PRJNA722386 and GSE172187) [86] was mapped aligned to the *R. irregularis* reference genome (GCF_026210795.1) using STAR with the option ‐‐outFilterMultimapNmax 20, counts were generated with featureCounts, differential gene expression analysis was performed using DESeq2 [87], and scaled normalised counts were plotted using the pheatmap package (version 1.0.12, https://rdrr.io/cran/pheatmap/).

### Spore germination assay

240,000 spores of *R. irregularis* DAOM197198 purchased from Premier Tech were rinsed three times with sterile water, then drained using 40-µM cell strainers and dried using sterile whatman paper. Spores were resuspended in sterile water, and 12 equal amounts were split into a 12-well tissue culture plate at a final concentration of 10,000 spores/mL (20,000 spores/well) using a cut-open P1000 tip. The plate was incubated at 28°C in the dark for 4 hours before collecting the day 0 replicates, which were individually strained and dried as described above, flash-frozen in liquid nitrogen and stored at -80°C. The plate was incubated at 28°C in the dark for collection of the day 7, 14 and 21 replicates.

### Microscopy

For *in planta* microscopy, *Oryza sativa* ssp. *japonica* cv. Nipponbare roots were harvested at 6 weeks post-inoculation (wpi) with *R. irregularis* DAOM197198 and segments (0.3–0.5 cm in length) were fixed in paraformaldehyde (PFA) and embedded in paraffin. Sections of 10 µm were adhered to slides, stained with WGA-AF488 and counterstained with DAPI to visualize both plant and fungal nuclei. Fluorescence imaging was performed using a Celldiscoverer 7 (Carl Zeiss Microscopy GmbH)) microscope with 25x water-immersion objective and automated focus finding via custom commands. The images (.czi) from each of the acquired channels for DAPI and WGA-AF488 were flattened by maximum Z-projection and the coherent regions of interest were stitched together to obtain respective panorama image files (.tiff) in Zen Blue software (Carl Zeiss Microscopy GmbH). For microscopy of *R. irregularis*, spores were germinated for 7 days as described above, stained with DAPI and mounted on fresh 2% agarose pads with 10 µL of water. Fluorescence live imaging was performed with a Leica DM6B. To quantify paired and non-paired nuclei, a box of 7.8×4.2 µm was used as a delimitation guide. Two clearly distinguishable DAPI foci that fit inside the box were counted as paired, and all others were counted as non-paired.

### DNA extraction and qPCR

Liquid nitrogen-frozen *R. irregularis* spore samples were homogenized using a mixer mill MM 400 and 25-mL grinding jars (Retsch), pre-chilled in liquid nitrogen and shaking at 25 shakes/sec for 20 sec. Samples were transferred to 50mL Falcons containing 17.5mL lysis buffer (100 mM tris-HCl, pH 8.0, 20 mM EDTA, 0.5 M NaCl, and 1% SDS) and 10uL RNAse (1000 U/ml, Thermo Fisher EN0541), and mixed well by vortexing. In order to control for DNA extraction, 7.8 pg of the spike-in plasmid pUC19 was added to each lysate, representing an approximate 1:1 ratio of plasmid:*R*.*irregularis* genome copy number. Lysates were incubated at room temperature for 30 minutes with mixing by inversion every 5 minutes. 200uL Proteinase K (800U/ml, NEB P81072) was added to each sample and incubated at room termerature for 30 minutes with mixing by inversion every 5 minutes. Samples were cooled in ice before adding 3.5 mL (0.2V) of KAc (5M pH 7.5), mixed by inversion, incubated on ice for 5 minutes, then spun at 4°C and 5000g for 12 minutes. Supernatants were transferred to fresh tubes containing 17.5 mL (1V) (P/C/I) and mix by inversion for 2 minutes, spun at 4°C and 4000g for 10 minutes. DNA was precipitated using 0.1V NaAc (3M pH 5.2) and 1V isopropanol for 10 minutes at room temperature. After centrifugation at 4 °C and 8000g for 30 mins, pellets were washed twice with 70% ethanol, air-dried and resuspended in Tris buffer (10mM pH 8.5). qPCRs were performed using the PowerTrack SYBR Green Master Mix (ThermoFisher), as described by the manufacturer. The primers M13_F 5’-GTAAAACGACGGCCAG-3’ and M13_R 5’-CAGGAAACAGCTATGAC-3’, Chr3_F 5’-GAAGTTCACGAGCTGCTTTATTAG-3’ and Chr3_R 5’-GTGAATCACATGCCATTTAAGGT-3’, Chr5_F 5’-TGTCTGTCCATTGCAGTGTC-3’ and Chr5_R 5’-GCGGTCCTTTAGTAGCCATATC-3’, Mito_F 5’-ATCCGCCTGCAGAACTAATC-3’ and Mito_R 5’-GGCTTCGAGCCTGTCTTATATC-3’ were used to amplify the spike-in plasmid pUC19, and *R. irregularis* chromosome 3, chromosome 5 and mitochondrial chromosome respectively.

## Data Availability

Genomic datasets used in this manuscript are publicly available on NCBI and accessions are listed in Table S1. The orthologous group dataset generated is available at https://doi.org/10.5281/zenodo.15428617.

## Acknowledgements

We are grateful to members of the Hayashi and Paszkowski groups for discussions, and to Hajk-Georg Drost, Michael Boemo, Sebastian Schornack, Navin Ramakrishna and Miguel Vasconcelos Almeida for critical reading and suggestions.

## Author contributions

Conceptualisation, investigation, data curation, formal analysis, visualisation, supervision, project administration, writing original draft: A.D.

Investigation, Writing – Review & Editing: all authors contributed.

